## Supplementary Methods for "DENcode: A model for haplotype-informed transmission probability of dengue virus"

Reconstruction of haplotype sequences and hotspot identification

The novel work presented in this paper includes DENcode probabilistic framework and its validation. The data used to validate the model had been generated (genetic data) or collected (clinical data) as part of previously published work and for completeness the process of data generation is briefly reproduced below (citing the corresponding prior publications).

Clinical data: For information on CDS including the design, aims and types of data collected please refer to previous publications (20, 23, 44, 45).

DENV genomic data: From CDS patient plasma samples, the near full length DENV genomes were extracted and amplified by a nested PCR as previously described (24). The product was cleaned by magnetic beads (Ampure®, Beckman Coulter, CA, USA), visualised on a 0.8% agarose gel and quantified (Quant-iT Picogreen® assay, ThermoFisher, USA, catalogue number: P7589). Deep sequencing with Oxford Nanopore Technology (ONT) was carried out by an external service provider, Kinghorn Centre for Clinical Genomics, Sydney, Australia using PromethION flow cells (FLO-PRO114M) with native barcoding (SQK-NBD114-96) with multiplexing between 24 – 48 samples in a single run. The base called data were filtered for high quality reads and demultiplexed with Dorado (7.3.11+0112dde09) to generate an output file in FASTq format.

The FASTq files were iteratively mapped on to a serotype specific reference sequence in Minimap2(46) implemented within Geneious Prime (version 2024, BioMatters Geneious, New Zealand) to generate consensus sequences for each sample as described previously (16). The haplotypes per patient were reconstructed using the corresponding FASTq file and the autologous consensus sequence with Nano-Q , a previously published bioinformatics pipeline that parses haplotype sequences and their relative abundances (47). While the dengue consensus sequences were sequenced from 345 out of 523 samples, only 104 samples had enough sequencing depth for successful haplotype reconstruction(16).

Certain amino acid loci mutate rapidly due to diversifying selection. To prevent these hypervariable sites from dominating distance estimates, we applied a hotspot-adjusted patristic distance. We had previously identified hotspots in this CDS viral genomic dataset both at consensus level and at haplotype levels(16). This prior analysis defined mutation hotspots as amino acid positions with significantly high Shannon Entropy (>mean + 2 standard deviations) and undergoing diversifying selection as identified by any one of the following algorithms: Fixed Effects Likelihood (FEL), A Fast, Unconstrained Bayesian AppRoximation for Inferring Selection (FUBAR), Mixed Effects Model of Evolution (MEME).
