## Supplementary Data for "DENcode: A model for haplotype-informed transmission probability of dengue virus"

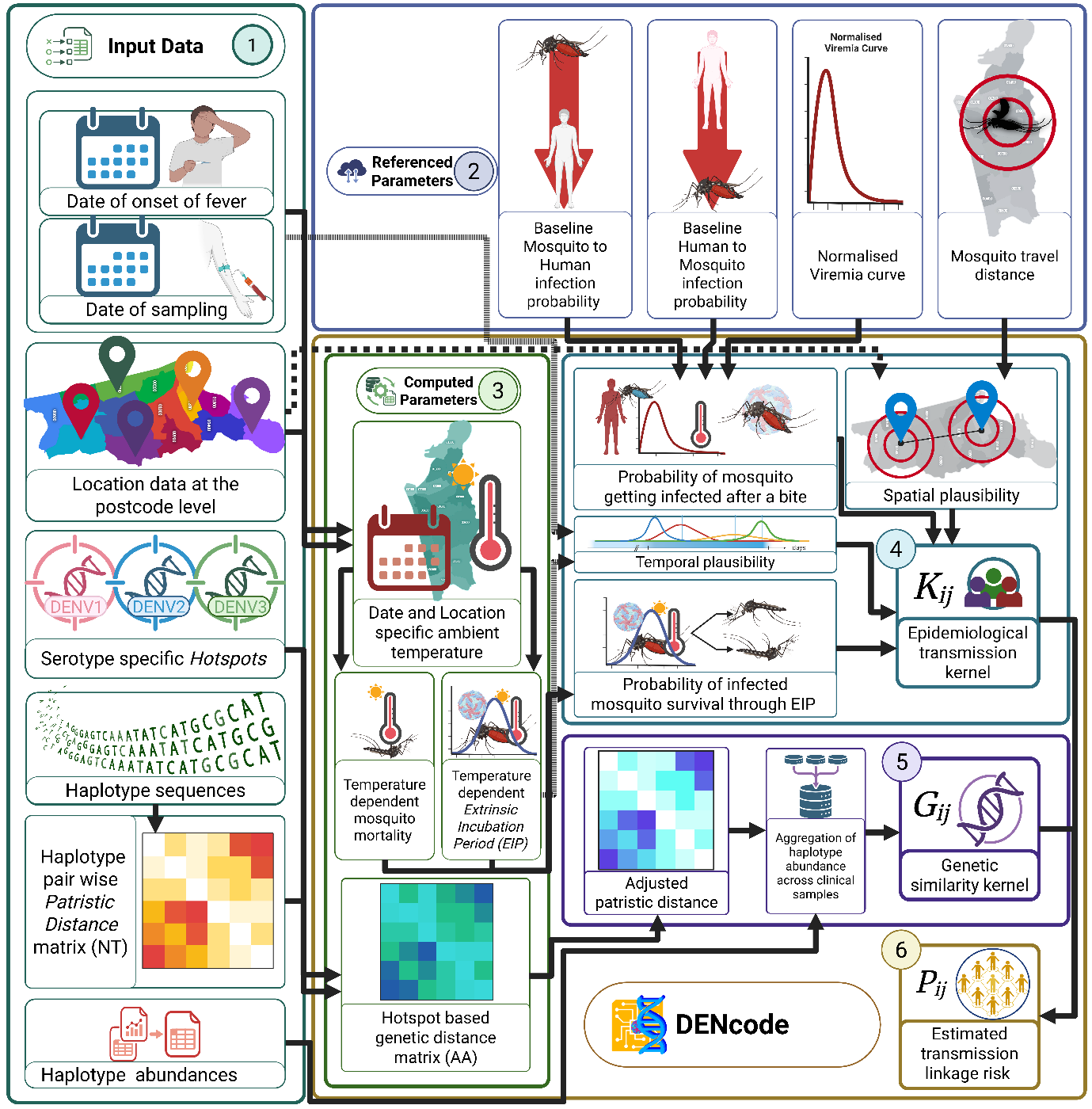

S Fig 1. Process flow diagram of DENcode. Input data (1) include epidemiological dates, geocoded locations, serotype-specific hotspot data, and within-host haplotype sequences. Referenced parameters (2) define baseline transmission probabilities and biological constraints. Computed parameters (3) integrate temperature-dependent vector dynamics and spatial genetic structure. The epidemiological kernel $\boldsymbol{K}_{\boldsymbol{ij}}$ (4) combines spatial plausibility, temporal compatibility, and extrinsic incubation period survival probability. The genetic kernel $\boldsymbol{G}_{\boldsymbol{ij}}$ (5) aggregates haplotype-level genetic distances. Finally, DENcode (6) estimates pairwise transmission probability $\boldsymbol{P}_{\boldsymbol{ij}}$by integrating epidemiological and genetic kernels. Framework uses complete haplotype sequences to capture within-host diversity, enabling high-resolution outbreak investigation.

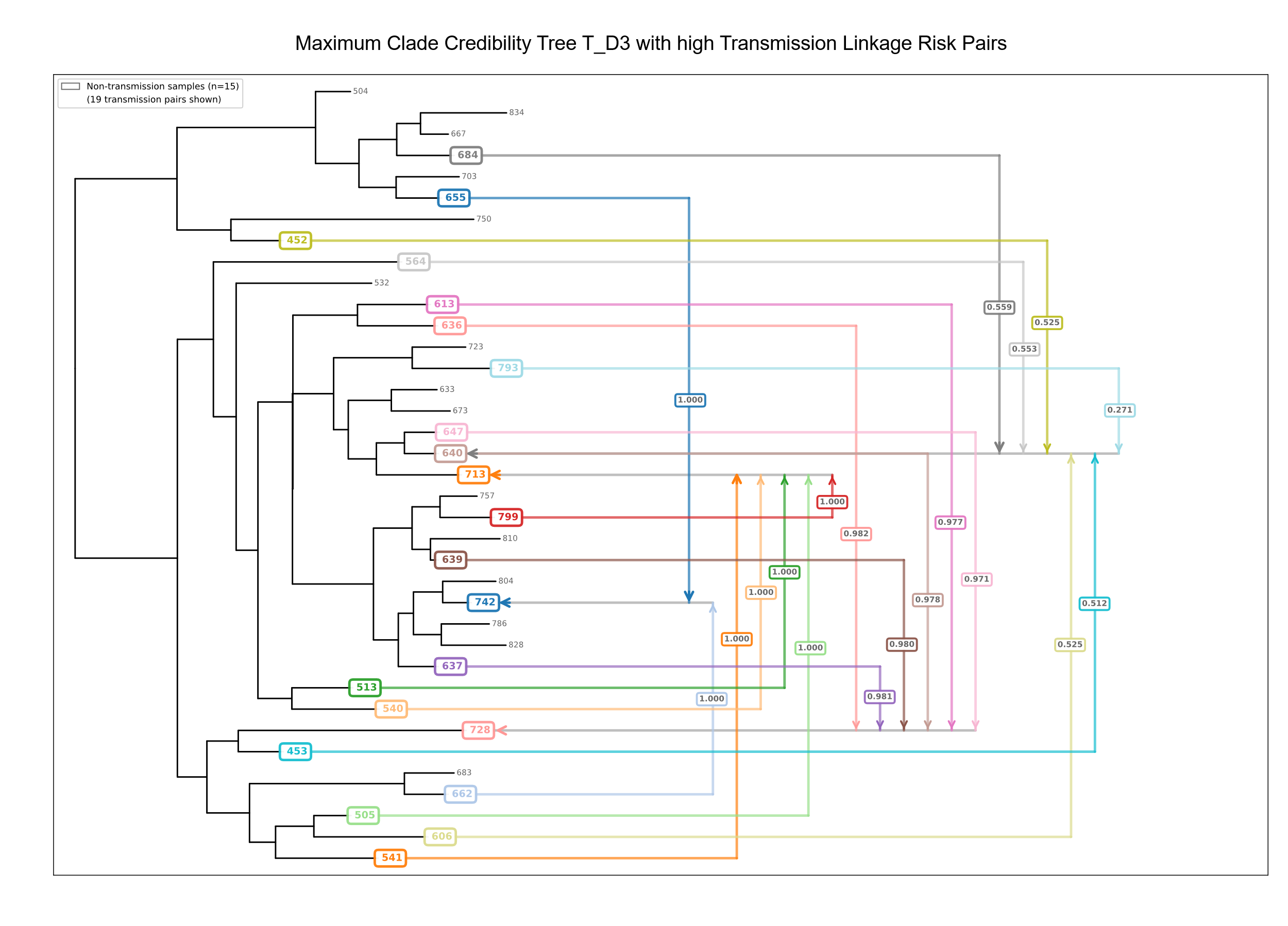

S Fig 2. Maximum Clade Credibility tree generated from cluster T(23) identified from the DENV3 samples of the Colombo Dengue Study highlighted with high transmission linkage risk pairs. The specific high probability linkages are marked by coloured arrows, and the transmission linkage risk is shown within each arrow.

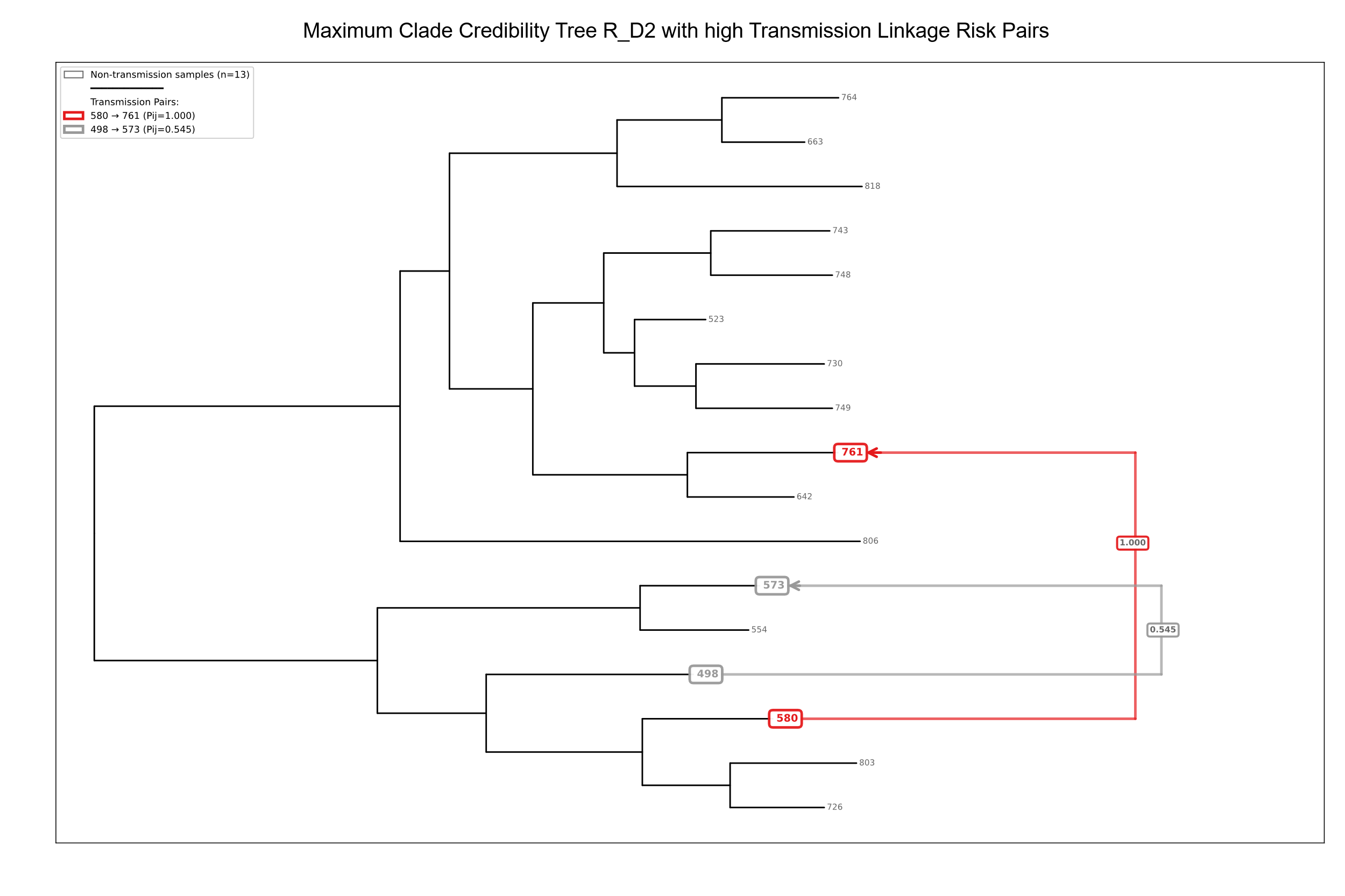

S Fig 3. Maximum Clade Credibility tree generated from cluster R (23) identified from the DENV2 samples of the Colombo Dengue Study highlighted with high transmission linkage risk pairs. The specific high probability linkages are marked by coloured arrows, and the transmission linkage risk is shown within each arrow.

**S table 1. Simulated directionality with the date of onset of fever**

| Serotype | Threshold | Total Edges | Edges With Dates | Concordant Edges | Concordant Percentage | Median Time Lag  (Days) |
| --- | --- | --- | --- | --- | --- | --- |
| D1 | ≥0.5 | 5 | 2 | 2 | 100 | 295.16 |
| D1 | ≥0.1 | 6 | 2 | 2 | 100 | 295.16 |
| D2 | ≥0.5 | 25 | 25 | 25 | 100 | 465.71 |
| D2 | ≥0.1 | 87 | 77 | 77 | 100 | 437.05 |
| D3 | ≥0.5 | 19 | 15 | 15 | 100 | 43.29 |
| D3 | ≥0.1 | 41 | 29 | 29 | 100 | 99.88 |

**S Table 2. Summary of statistical comparison of transmission pairs and non-transmission branches in the two-cluster specific MCC BEAST trees.**

| Tree | Serotype | Samples | Transmission Pairs | Mean (SD)  Transmission Distance | Mean (SD)  Non-Transmission Pair Distance | Mann Whitney U test P value | Significance |
| --- | --- | --- | --- | --- | --- | --- | --- |
| R_D2 | D2 | 17 | 2 | 3.255 (1.112) | 3.084 (1.295) | 0.568 | No |
| T_D3 | D3 | 37 | 19 | 1.663 (0.436) | 1.648 (0.673) | 0.554 | No |
